# Associations between fluid biomarkers and PET imaging ([^11^C]UCB-J) of synaptic pathology in Alzheimer’s disease

**DOI:** 10.1101/2025.03.31.646290

**Authors:** Johanna Nilsson, Adam P. Mecca, Nicholas J. Ashton, Elaheh Salardini, Ryan S. O’Dell, Richard E. Carson, Andrea L. Benedet, Kaj Blennow, Henrik Zetterberg, Christopher H. van Dyck, Ann Brinkmalm

## Abstract

**INTRODUCTION:** Positron Emission Tomography (PET) imaging with ligands for synaptic vesicle glycoprotein 2A (SV2A) has emerged as a promising methodology for measuring synaptic density in Alzheimer’s disease (AD). We investigate the relationship between SV2A PET and CSF synaptic protein changes of AD patients.

**METHOD:** Twenty-one participants with early AD and 7 cognitively normal (CN) individuals underwent [^11^C]UCB-J PET. We used mass spectrometry to measure a panel of synaptic proteins in CSF.

**RESULTS:** In the AD group, higher levels of syntaxin-7 and PEBP-1 were associated with lower global synaptic density. In the total sample, lower global synaptic density was associated with higher levels of AP2B1, neurogranin, γ-synuclein, GDI-1, PEBP-1, syntaxin-1B, and syntaxin-7 but not with the levels of the neuronal pentraxins or 14-3-3 zeta/delta.

**CONCLUSION:** Reductions of synaptic density found in AD compared to CN participants using [^11^C]UCB-J PET were observed to be associated with CSF biomarker levels of synaptic proteins.

## BACKGROUND

Synaptic dysfunction and degeneration are an early and significant part of the pathology occurring in Alzheimer’s disease (AD) as well as in other neurodegenerative diseases [1]. Postmortem brain studies have described a widespread reduction in synapse numbers in AD in comparison to healthy individuals and this synaptic loss is the major structural correlate of cognitive decline, even more so than amyloid-beta (Aβ) plaque pathology [2]. These findings have contributed to an increasing interest in the development and implementation of synaptic biomarkers in diagnostics as well as mechanistic insight into synaptic pathology.

Already 30 years ago, the first studies emerged verifying the presence of synaptic proteins in the cerebrospinal fluid (CSF). Since then, several methods for the quantification of a wide range of synaptic proteins in living patients have been developed as synaptic biomarkers [3]. One of the most established synaptic biomarkers is neurogranin, a postsynaptic protein that is specifically increased in AD with changes at early stages of cognitive impairment [4] and even before symptom onset [5]. Other synaptic proteins including SNAP-25, synaptotagmin-1, and GAP43 increase along with biomarkers of cerebral amyloid pathology in symptomatic and asymptomatic disease, and are considered AD specific (*e*.*g*.,) [6-8]. Interestingly, some synaptic proteins are lower in CSF samples of participants with neurodegenerative disease. For example, neuronal pentraxins have decreased levels across symptomatic neurodegenerative diseases, including AD, compared to healthy individuals and the levels of pentraxins strongly correlate with cognitive decline [9-11]. Emerging evidence thus points to the fact that synaptic pathology mechanisms are more complex than earlier believed and that all synaptic proteins may not represent the same pathological pathways.

Recent progress in synaptic positron emission tomography (PET) imaging has also allowed for the evaluation of synaptic alterations *in vivo* [1, 12]. Utilizing [^11^C]UCB-J, a PET tracer that binds the synaptic vesicle glycoprotein 2A (SV2A) which is expressed in nearly all synapses, the synaptic density of patients with AD was found to be reduced in the medial temporal and neocortical brain regions compared with healthy controls [12, 13]. SV2A PET has been used to characterize the patterns of synaptic loss due to neurodegenerative disease and early studies have described complex relationships with core AD biomarkers such as greater synaptic los in areas of tau deposision [14], significant correlations between synaptic density and metabolism measured with [^18^F]FDG [15], and stage dependent correlations between hippocampal synaptic density and global amyloid measured with [^11^C]PiB [16]. In addition, SV2A PET has been used to demonstrated significant synaptic loss in dementia with Lewy bodies [17], frontotemporal dementia [18], Parkinson’s disease [19], and Huntington’s disease [20]. Thus, both imaging and CSF protein biomarkers are altered in AD and other neurodegenerative disorders and these changes are likely reflect synaptic pathology. However, the meaning of synaptic protein concentration changes measured in vivo with PET or CSF assays remains unclear. These changes may reflect altered clearance or altered protein production and secretion into the CSF due to synapse degeneration or changes in synaptic activity [21].

In the present study, we evaluate the relationship between SV2A PET and CSF synaptic protein changes of AD patients to gain insight into AD synaptic pathology and the meaning of synaptic protein biomarkers. To do so, we have measured a panel of synaptic proteins in a small cohort of AD patients and control participants who also have undergone SV2A PET.

## METHOD

### Study design and population

The requirement of participation included fulfilling the diagnostic criteria for amnestic mild cognitive impairment (MCI, n=5) [22] or probable dementia due to AD (n=16) [23]. AD and MCI participants were additionally required to have a positive PET scan with [^11^C]Pittsburgh Compound B ([^11^C]PiB), a Mini-Mental State Examination (MMSE) score <26 and 24-30, respectively, and a Clinical Dementia Rating (CDR) score of 0.5-1.0 and 0.5, respectively. Cognitively normal participants (CN, n=7) were required to have a negative [^11^C]PiB PET scan, MMSE score <26, and a CDR score of 0. All participants gave written informed consent approved by the Yale University Human Investigation Committee.

### Brain imaging

To define regions of interest (ROIs) and perform partial volume correction (PVC) using the iterative Yang approach [24], T1-weighted magnetic resonance imaging (MRI) was performed. FreeSurfer [version 6.0] was used for volumetric segmentation and cortical reconstruction. A high-resolution research tomograph (207 slices, resolution <3 mm, full width at half maximum (FWHM)) was used for the PET scans with event-by-event motion correction. Dynamic [^11^C]PiB and [^11^C]UCB-J scans 60 minutes, following administration of a bolus of up to 555 and 740 MBq of tracer, respectively. As previously described, SRTM2 (cerebellum reference region) was used with dynamic scan data from 0 to 60 min to generate parametric images of [^11^C]PiB *BP*_ND_ and [^11^C]UCB□J *DVR* [24].

### LC-MS analysis

CSF samples were collected in polypropylene tubes by lumbar puncture, centrifugated (2200 x g for 10 min, 20^°^C), and stored at -80^°^C. Eleven synaptic proteins were analyzed in the synaptic panel analysis; neurogranin, γ-synuclein, the activating protein 2 subunit complex beta (AP2B1), rab GDP dissociation inhibitor alpha (GDI-1), phosphatidylethanolamine-binding protein 1 (PEBP-1), 14-3-3 ζ/δ, syntaxin-1B, syntaxin-7, and the neuronal pentraxins (-1 (NPTX1), -2 (NPTX2), and the receptor (NPTXR)). The sample preparation of 100 μL of CSF samples entailed the addition of internal standard (mix of stable-isotope-labeled peptides, 25 μL, 0.032 pmol/μL), reduction, alkylation, tryptic digestion, and solid-phase extraction, (for thorough protocol, refer to [25]). Quantification of the synaptic proteins was performed on a micro-high-performance liquid chromatography-mass-spectrometry system (6495 Triple Quadrupole LC/MS system, Agilent Technologies) equipped with a Hypersil Gold reversed-phase C18 column (dim.=100×2.1 mm, particle size=1.9 μm, Thermo Fisher Scientific), for detailed settings see Suppl. Table 1. Assay performance was evaluated during the run by injections at regular intervals of a quality control sample.

**Table 1.** Demographics and biomarker characteristics.

|  | CN (N=7) | AD (N=21) | P-value |
| --- | --- | --- | --- |
| Sex, n (Male, %) | 5 (71.4%) | 10 (47.6%) | 0.274 |
| Age, y | 73.4 (7.3) | 69.1 (8.7) | 0.246 |
| MMSE score | 28.9 (1.35) | 23.0 (2.9) | <0.0001**** |
| A$\beta_{42/40}$ ratio | 0.086 (0.008) | 0.041 (0.009) | <0.0001**** |
| P-tau<sub>181</sub>, ng/L | 27.8 (11.2) | 123.0 (75.6) | 0.0030** |
| T-tau, ng/L | 291 (86) | 783 (449) | 0.0085** |
| NFL, ng/L | 1112 (592) | 1702 (820) | 0.092 |
| AP2B1, fmol/μL | 0.59 (0.30) | 0.96 (0.4) | 0.057 |
| γ-synuclein, fmol/μL | 0.41 (0.15) | 0.61 (0.19) | 0.022* |
| Neurogranin, fmol/μL | 0.027 (0.019) | 0.060 (0.032) | 0.017* |
| NPTXR, fmol/μL | 2.32 (1.36) | 2.93 (1.11) | 0.247 |
| NPTX1, fmol/μL | 1.36 (1.11) | 1.72 (0.80) | 0.357 |
| NPTX2, fmol/μL | 0.72 (0.37) | 0.94 (0.33) | 0.162 |
| GDI-1, fmol/μL | 0.17 (0.063) | 0.28 (0.089) | 0.0059** |
| PEBP-1, fmol/μL | 14.9 (6.3) | 24.4 (7.2) | 0.0047** |
| 14-3-3 $\zeta/\delta$, fmol/μL | 1.24 (0.60) | 2.26 (0.69) | 0.0018** |
| Syntaxin-1B, fmol/μL | 0.10 (0.060) | 0.18 (0.079) | 0.018* |
| Syntaxin-7, fmol/μL | 0.019 (0.007) | 0.025 (0.008) | 0.089 |

Notes: Data presented as mean (standard deviation). Analysis of variance (ANOVA) and chi-
square goodness of fit test were used to compare continuous and categorical variables
between groups. P-values: \* $P \leq 0.05$ , \*\* $P \leq 0.01$ , \*\*\* $P \leq 0.001$ and \*\*\*\* $P \leq 0.0001$ .
Abbreviations: AP2B1, adaptor-related protein complex 2 subunit beta 1; AD, Alzheimer's
disease; A $\beta_{42/40}$ , Amyloid-β 42/40; CN, Cognitively Normal; MMSE, Mini-Mental State Exam;
NFL, neurofilament light; NPTX, neuronal pentraxin; PEBP-1, phosphatidylethanolamine-
binding protein 1; P-tau<sub>181</sub>, tau phosphorylated at Thr181; GDI-1, rab GDP dissociation
inhibitor alpha; T-tau, total tau.

### Data processing and statistical analysis

Skyline 20.1 (MacCoss Lab Software) was utilized for peak inspection and adjustment of the chromatographic spectra and R software was used for the statistical analysis. For the proteins for which more than one peptide was analyzed, the peptide with the best repeatability (lowest CV), was chosen for the statistical analysis (Suppl. Table 2). The group comparisons of the demographic characteristics and biomarkers were evaluated by analysis of variance (ANOVA) and chi-square goodness of fit test for continuous and categorical variables, respectively. Associations between synaptic density and CSF biomarker levels were explored with Pearson’s rank correlation analysis and brain maps for visualization were created by setting each brain region’s voxels uniformly to the calculated effect size (Pearson’s *r*).

## RESULTS

### Participant characteristics and biomarker levels

The study sample was well-balanced regarding age and sex and the AD participants had both typical clinical characteristics (MMSE=23.0±2.9) and core CSF biomarker levels of total tau (t-tau), tau phosphorylated at Thr181 (p-tau_181_), amyloid-β 42/40 (Aβ_42/40_) (Table). Out of the 11 synaptic proteins quantified 6 proteins (neurogranin, γ-synuclein, GDI-1, PEBP-1, 14-3-3 ζ/δ, and syntaxin-1B) showed significantly higher protein levels (p<0.05) in the AD group compared to CN.

### Association between synaptic density and synaptic proteins

The primary analysis investigated the association between global synaptic density (*DVR*) in a composite of AD-affected regions [24] and synaptic protein levels in the AD group. We also examined the same associations in the the CN group and the total sample (Suppl. Table 3). In the AD group, significantly higher levels of syntaxin-7 (*r*=-0.49, p-value=0.024) and PEBP-1 (*r*=-0.47, p-value=0.033) were associated with lower global synaptic density. In the total sample, lower global synaptic density was significantly associated with higher levels of AP2B1, neurogranin, γ-synuclein, GDI-1, PEBP-1, syntaxin-1B, and syntaxin-7 (*r*=-0.40 to -0.54, p-value<0.05) but not with the levels of the neuronal pentraxins or 14-3-3 zeta/delta. In the CN group, there were no significant associations between any proteins and global synaptic density. When PVC was performed on PET data, in the AD group, only significantly higher levels of syntaxin-7 (*r*=-0.51, p-value<0.05) were associated with lower PVC global synaptic density. In the total sample, lower global synaptic density was significantly associated with higher levels of AP2B1, GDI-1, and PEBP-1 (*r*=-0.40 to -0.44, p-value<0.05). In the CN group, there were no significant associations between any proteins and PVC global synaptic density. When additional analyses assessed the association between CSF protein biomarkers and synaptic density in all brain regions, negative associations were found across many parietal, temporal, prefrontal, and occipital cortical regions (data not shown). The associations varied between biomarkers as seen by syntaxin-7, which showed the strongest associations with synaptic density in different brain regions, and neuronal pentraxin-2, which showed no associations with synaptic density as measured (Figure). Notably, in the context of AD where the hippocampus is known to be an area of early synaptic loss, there were no significant associations between any synaptic proteins and hippocampal synaptic density in either the separate groups or total sample (Figure and Suppl. Table 4).

**Figure.**
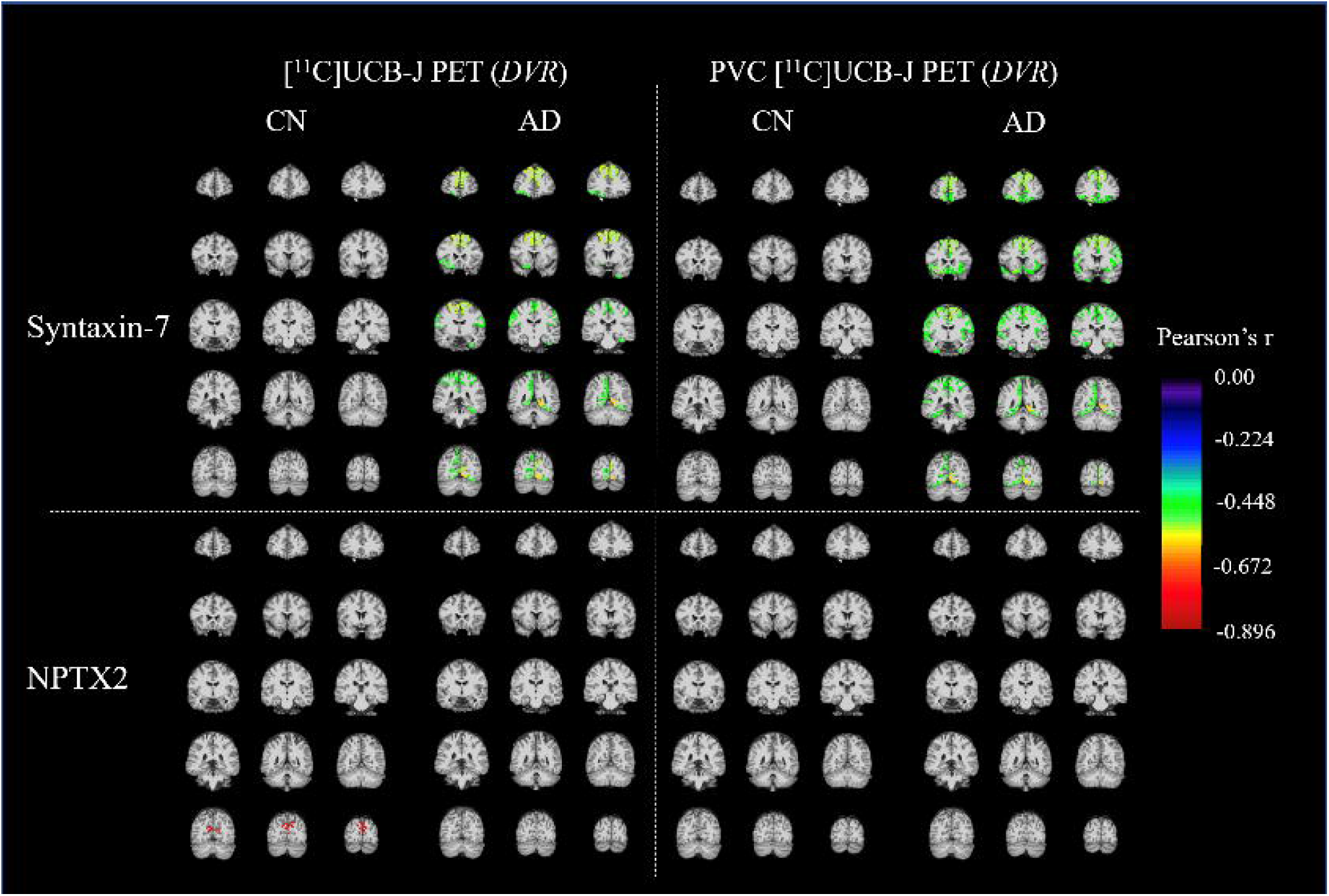
Regional associations (Pearson’s *r*) between cerebrospinal fluid syntaxin-7 or neuronal pentraxin-2 (NPTX2) and synaptic density (*DVR*) determined by [^11^C]UCB-J PET in Alzheimer’s disease (AD) and cognitively normal (CN) participant groups. Analysis was performed with and without partial volume correction (PVC) of PET data.

## DISCUSSION

In this study, we examined the relationship between potential biomarkers of synaptic pathology and synaptic density using a mass spectrometric synaptic protein panel and [^11^C]UCB-J PET. In the AD group, the most robust global associations were seen for syntaxin-7 and PEBP-1. While in the total sample, associations were found for several of the synaptic proteins, including the established synaptic biomarker neurogranin. Interestingly, no associations were found for the neuronal pentraxins, synaptic proteins that are emerging as interesting biomarkers in neurodegenerative diseases.

Syntaxin-7 is involved in vesicle endocytosis at the synapse, as a SNARE protein, and thus mediation of endocytic trafficking [26]. Endocytic impairment has been implicated to be a feature of many neurodegenerative diseases, not the least in AD [27]. Studies of syntaxin-7 as a potential biomarker have however shown no changes in the CSF of AD patients or other diseases compared to controls [9, 25]. We also observed a non-significant difference between AD patients when compared to controls in the current study. However, syntaxin-7 was shown to have the strongest association with synaptic density as quantified by [^11^C]UCB-J PET within AD patients. Since SV2A, the target for [^11^C]UCB-J, and syntaxin-7 are both synaptic vesicle proteins the stronger associations between synaptic density and syntaxin-7 compared to the other synaptic proteins might be attributed to the proximity in function between the two. PEBP-1 was also associated with [^11^C]UCB-J PET in AD patients, but displayed higher levels in the AD compared to the CN group. PEBP-1 is a regulatory protein with modulatory roles in several protein kinase signaling cascades as well as the precursor to the hippocampal cholinergic neurostimulating peptide (HCNP) implicated in the induction of acetylcholine synthesis and enhancement of glutamatergic activity [28, 29].

The neuronal pentraxins are three synaptic proteins, two secreted glycoproteins (NPTX1 and NPTX2), and their plasma membrane-anchored receptor (NPTXR), which have important roles in synaptic function and plasticity [30]. By associating and forming heteromultimers the neuronal pentraxins are involved in the recruitment and localization of neurotransmitter receptors to the postsynaptic membrane during receptor exocytosis. As previously mentioned, pentraxins have been recognized as potential synaptic biomarkers with decreased levels across neurodegenerative diseases compared to healthy individuals and whose levels strongly correlate with cognitive decline [9, 10, 31-36]. However, in the current study, neuronal pentraxins levels in the CSF did not differ between AD and CN groups, and did not correlate with reduced synaptic density in AD. The cause of this inconsistency is not clear, but differences in AD pathologic stage or other participant characteristics between sample studies could be contributing. In combination with the observation that the levels of the neuronal pentraxins in the CSF of AD patients differ in the directionality of change (decreased instead of increased) compared with most other synaptic proteins might further implicate that the neuronal pentraxins levels in CSF are indicative of a unique synaptic pathologic processes occurring in AD. Further studies are needed to confirm these findings and to investigate what possible mechanisms might underlie the difference between synaptic protein alterations in AD.

The main limitation of this study is the modest sample size leading to limited power of the study to detect associations. This might have especially affected the control group where no associations were seen, nevertheless, all scatter plots (data not shown) were visually inspected and no convincing associations were observed. However, we are encouraged that several proteins including syntaxin-7 and PEBP-1 demonstrated associations with [^11^C]UCB-J PET in AD. Future studies with larger sample sizes and a broader range of synaptic proteins may permit a better understanding of the relationship between altered synaptic density fluid synaptic biomarkers in AD.

## CONCLUSIONS

We observed that the reductions of synaptic density found in AD compared to CN participants using [^11^C]UCB-J PET are associated with biomarker levels of synaptic proteins.

## ABBREVIATIONS

AP2B1: adaptor related protein complex 2 subunit beta 1
AD: Alzheimer’s disease
Aβ: amyloid-β Aβ_42/40,_ Amyloid-β 42/40
CDR: Clinical Dementia Rating
CSF: cerebrospinal fluid
CV: coefficient of variation
MMSE: Mini-Mental State Examination
NPTX: neuronal pentraxin
PEBP-1: phosphatidylethanolamine-binding protein 1
PET: positron emission tomography
P-tau_181_: tau phosphorylated at Thr181
GDI-1: rab GDP dissociation inhibitor alpha
ROI: region of interest
PVC: partial volume correction; *r*, Pearson’s rank correlation coefficient
SV2A: synaptic vesicle glycoprotein 2A
T-tau: total tau.

## Ethics approval and consent to participate

## Consent for publication

Not applicable

## Availability of data and materials

Derived data supporting the findings of this study are available from the corresponding author on request, providing data transfer is in agreement with the participating centre national legislation and institutional review centre.

## Competing interests

HZ has served at scientific advisory boards and/or as a consultant for Abbvie, Alector, Annexon, AZTherapies, CogRx, Denali, Eisai, Nervgen, Pinteon Therapeutics, Red Abbey Labs, Roche, Samumed, Siemens Healthineers, Triplet Therapeutics, and Wave, has given lectures in symposia sponsored by Cellectricon, Fujirebio, Alzecure and Biogen, and is a co-founder of Brain Biomarker Solutions in Gothenburg AB (BBS), which is a part of the GU Ventures Incubator Program. KB has served as a consultant, at advisory boards, or at data monitoring committees for Abcam, Axon, Biogen, JOMDD/Shimadzu. Julius Clinical, Lilly, MagQu, Novartis, Roche Diagnostics, and Siemens Healthineers, and is a co-founder of Brain Biomarker Solutions in Gothenburg AB (BBS), which is a part of the GU Ventures Incubator Program. APM reports grants for clinical trials from Genentech, Eli Lilly, and Janssen Pharmaceuticals outside the submitted work. CHvD reports consulting fees from Kyowa Kirin, Roche, Merck, Eli Lilly, and Janssen and grants for clinical trials from Biogen, Novartis, Eli Lilly, Merck, Eisai, Janssen, Roche, Genentech, Toyama, and Biohaven, outside the submitted work. The other authors declare no conflict of interest.

## Funding

This work was supported by the Eivind and Elsa K: Son Sylvan Foundation, the Foundation for Gamla Tjänarinnor, and Demensfonden. HZ is a Wallenberg Scholar supported by grants from the Swedish Research Council (#2018-02532), the European Research Council (#681712), Swedish State Support for Clinical Research (#ALFGBG-720931), the Alzheimer Drug Discovery Foundation (ADDF), USA (#201809-2016862), the AD Strategic Fund and the Alzheimer’s Association (#ADSF-21-831376-C, #ADSF-21-831381-C and #ADSF-21-831377-C), the Olav Thon Foundation, the Erling-Persson Family Foundation, Stiftelsen för Gamla Tjänarinnor, Hjärnfonden, Sweden (#FO2019-0228), the European Union’s Horizon 2020 research and innovation programme under the Marie Skłodowska-Curie grant agreement No 860197 (MIRIADE), and the UK Dementia Research Institute at UCL. KB is supported by the Swedish Research Council (#2017-00915), the Alzheimer Drug Discovery Foundation (ADDF), USA (#RDAPB-201809-2016615), the Swedish Alzheimer Foundation (#AF-742881), Hjärnfonden, Sweden (#FO2017-0243), the Swedish state under the agreement between the Swedish government and the County Councils, the ALF-agreement (#ALFGBG-715986), the European Union Joint Program for Neurodegenerative Disorders (JPND2019-466-236), the National Institute of Health (NIH), USA, (grant #1R01AG068398-01), and the Alzheimer’s Association 2021 Zenith Award (ZEN-21-848495). APM, RSO, REC and CHvD received support for this work from the National Institute on Aging (P30AG066508, P50AG047270, R01AG052560, R01AG062276).

Funders did not have any role in the study design, data collection or analysis and interpretation of the results or manuscript writing.

## Authors’ contributions

JN, NJA, and APM designed the study. JN performed the experiments. JN, NJA, ES, RSO and APM analysed the data and wrote the manuscript. AB, HZ, KB were responsible for supervision, conceptualization, and verification of the underlying data. APM, RSO, REC and CHvD provided the CSF samples of the clinical cohort. APM, RSO and CHvD participated in diagnosis of the patients and CSF samples collection. AB, ES, REC, CHvD, HZ, and KB contributed to the interpretation of the results and provided critical feedback of the manuscript. All authors have reviewed the manuscript.

## Acknowledgements

Not applicable

## REFERENCES

1. Camporesi, E., et al., Fluid Biomarkers for Synaptic Dysfunction and Loss. Biomarker Insights, 2020. 15: p. 1177271920950319.

2. Terry, R.D., et al., Physical basis of cognitive alterations in Alzheimer’s disease: synapse loss is the major correlate of cognitive impairment. Annals of Neurology: Official Journal of the American Neurological Association and the Child Neurology Society, 1991. 30(4): p. 572–580.

3. Davidsson, P., M. Puchades, and K. Blennow, Identification of synaptic vesicle, pre- and postsynaptic proteins in human cerebrospinal fluid using liquid-phase isoelectric focusing. Electrophoresis, 1999. 20(3): p. 431–437.

4. Mavroudis, I.A., et al., A meta-analysis on CSF neurogranin levels for the diagnosis of Alzheimer’s disease and mild cognitive impairment. Aging Clin Exp Res, 2019.

5. Milà-Alomà, M., et al., Amyloid beta, tau, synaptic, neurodegeneration, and glial biomarkers in the preclinical stage of the Alzheimer’s continuum. Alzheimer’s & Dementia, 2020.

6. Nilsson, J., et al., Quantification of SNAP-25 with mass spectrometry and Simoa: a method comparison in Alzheimer’s disease. Alzheimer’s Research & Therapy, 2022. 14(1): p. 1–10.

7. Milà-Alomà, M., et al., CSF synaptic biomarkers in the preclinical stage of Alzheimer disease and their association with MRI and PET: a cross-sectional study. Neurology, 2021. 97(21): p. e2065–e2078.

8. Öhrfelt, A., et al., Association of CSF GAP-43 With the Rate of Cognitive Decline and Progression to Dementia in Amyloid-Positive Individuals. Neurology, 2023. 100(3): p. e275–e285.

9. Nilsson, J., et al., Cerebrospinal fluid biomarker panel of synaptic dysfunction in Alzheimer’s disease and other neurodegenerative disorders. Alzheimer’s & Dementia, 2022.

10. Nilsson, J., et al., Cerebrospinal Fluid Biomarkers of Synaptic Dysfunction Are Altered in Parkinson’s Disease and Related Disorders. Movement Disorders, 2022.

11. Sogorb-Esteve, A., et al., Differential impairment of cerebrospinal fluid synaptic biomarkers in the genetic forms of frontotemporal dementia. Alzheimer’s research & therapy, 2022. 14(1): p. 1–12.

12. Chen, M.-K., et al., Assessing synaptic density in Alzheimer disease with synaptic vesicle glycoprotein 2A positron emission tomographic imaging. JAMA neurology, 2018. 75(10): p. 1215–1224.

13. Mecca, A.P., et al., In vivo measurement of widespread synaptic loss in Alzheimer’s disease with SV2A PET. Alzheimer’s & Dementia, 2020. 16(7): p. 974–982.

14. Mecca, A.P., et al., Association of entorhinal cortical tau deposition and hippocampal synaptic density in older individuals with normal cognition and early Alzheimer’s disease. Neurobiol Aging, 2022. 111: p. 44–53.

15. Chen, M.K., et al., Comparison of [(11)C]UCB-J and [(18)F]FDG PET in Alzheimer’s disease: A tracer kinetic modeling study. J Cereb Blood Flow Metab, 2021. 41(9): p. 2395–2409.

16. O’Dell, R.S., et al., Association of Abeta deposition and regional synaptic density in early Alzheimer’s disease: a PET imaging study with [(11)C]UCB-J. Alzheimers Res Ther, 2021. 13(1): p. 11.

17. Andersen, K.B., et al., Reduced synaptic density in patients with lewy body dementia: An [11C] UCB-J PET imaging study. Movement Disorders, 2021. 36(9): p. 2057–2065.

18. Salmon, E., et al., In vivo exploration of synaptic projections in frontotemporal dementia. Scientific Reports, 2021. 11(1): p. 16092.

19. Matuskey, D., et al., Synaptic changes in Parkinson disease assessed with in vivo imaging. Annals of neurology, 2020. 87(3): p. 329–338.

20. Delva, A., et al., Synaptic damage and its clinical correlates in people with early Huntington disease: a PET study. Neurology, 2022. 98(1): p. e83–e94.

21. Heurling, K., et al., Synaptic vesicle protein 2A as a potential biomarker in synaptopathies. Molecular and Cellular Neuroscience, 2019. 97: p. 34–42.

22. Albert, M.S., et al., The diagnosis of mild cognitive impairment due to Alzheimer’s disease: recommendations from the National Institute on Aging-Alzheimer’s Association workgroups on diagnostic guidelines for Alzheimer’s disease. Alzheimer’s & dementia, 2011. 7(3): p. 270–279.

23. McKhann, G.M., et al., The diagnosis of dementia due to Alzheimer’s disease: Recommendations from the National Institute on Aging-Alzheimer’s Association workgroups on diagnostic guidelines for Alzheimer’s disease. Alzheimer’s & dementia, 2011. 7(3): p. 263–269.

24. Mecca, A.P., et al., Synaptic density and cognitive performance in Alzheimer’s disease: A PET imaging study with [(11) C]UCB-J. Alzheimers Dement, 2022.

25. Nilsson, J., et al., Cerebrospinal fluid biomarker panel for synaptic dysfunction in Alzheimer’s disease. Alzheimer’s & Dementia: Diagnosis, Assessment & Disease Monitoring, 2021. 13(1): p. e12179.

26. Teng, F.Y.H., Y. Wang, and B.L. Tang, The syntaxins. Genome biology, 2001. 2(11): p. reviews3012. 1.

27. Root, J., et al., Lysosome dysfunction as a cause of neurodegenerative diseases: Lessons from frontotemporal dementia and amyotrophic lateral sclerosis. Neurobiology of disease, 2021. 154: p. 105360.

28. Yeung, K., et al., Mechanism of suppression of the Raf/MEK/extracellular signal-regulated kinase pathway by the raf kinase inhibitor protein. Molecular and cellular biology, 2000. 20(9): p. 3079–3085.

29. Madokoro, Y., et al., Reduced Cholinergic Activity in the Hippocampus of Hippocampal Cholinergic Neurostimulating Peptide Precursor Protein Knockout Mice. International journal of molecular sciences, 2019. 20(21): p. 5367.

30. Gómez de San José, N., et al., Neuronal pentraxins as biomarkers of synaptic activity: from physiological functions to pathological changes in neurodegeneration. Journal of Neural Transmission, 2021: p. 1–24.

31. Galasko, D., et al., Synaptic biomarkers in CSF aid in diagnosis, correlate with cognition and predict progression in MCI and Alzheimer’s disease. Alzheimers Dement (N Y), 2019. 5: p. 871–882.

32. Spellman, D.S., et al., Development and evaluation of a multiplexed mass spectrometry based assay for measuring candidate peptide biomarkers in Alzheimer’s Disease Neuroimaging Initiative (ADNI) CSF. Proteomics Clin Appl, 2015. 9(7-8): p. 715–31.

33. Swanson, A., A. Willette, and A.s.D.N. Initiative, Neuronal pentraxin 2 predicts medial temporal atrophy and memory decline across the Alzheimer’s disease spectrum. Brain, behavior, and immunity, 2016. 58: p. 201–208.

34. Begcevic, I., et al., Neuronal pentraxin receptor-1 is a new cerebrospinal fluid biomarker of Alzheimer’s disease progression. F1000Res, 2018. 7: p. 1012.

35. Lim, B., et al., Cerebrospinal fluid neuronal pentraxin receptor as a biomarker of long-term progression of Alzheimer’s disease: a 24-month follow-up study. Neurobiology of Aging, 2020.

36. Libiger, O., et al., Longitudinal CSF proteomics identifies NPTX2 as a prognostic biomarker of Alzheimer’s disease. Alzheimer’s & Dementia, 2021.

