## supplementary for "Associations between fluid biomarkers and PET imaging ([^11^C]UCB-J) of synaptic pathology in Alzheimer’s disease"

### **Supplementary Tables and Figures**

**Supplementary Table 1.** LC-MS/MS settings for the analysis of the synaptic protein panel.

|  | **Parameter** | **Setting** |
| --- | --- | --- |
| **LC** | Sample injection volume | 40 µL |
|  | Flow-rate | 0.3 mL/min |
|  | Gradient | Broken; 5–20%B (20 min), 20–35%B (7 min) |
|  | Total cycle time | 30 min |
|  | Mobile phase A | 0.1% formic acid in water (v/v) |
|  | Mobile phase B | 0.1% formic acid/84% acetonitrile in water (v/v) |
| **Electrospray** | Mode | Positive |
|  | Gas temperature | 220 °C |
|  | Gas flow | 15 L/min |
|  | Nebulizer pressure | 40 psi |
|  | Sheath gas temperature | 200 °C |
|  | Sheath gas flow | 11 L/min |
|  | Capillary voltage | 3500 V |
|  | Nozzle voltage | 500 V |
| **iFunnel** | Mode | Positive |
|  | High-pressure radio frequency | 200 V |
|  | Low-pressure radio frequency | 160 V |
| **MRM method** | Retention time window | 0.8 min |
|  | Collision energies | Individually optimized per transition |
|  | Cell accelerator voltage | Individually optimized per transition |

**Supplementary Table 2**. Calculations performed according to formulas from ISO 5725-2 section 7.4. Proteins/peptides marked as bold continued with for statistical analysis.

|  | QC | |
| --- | --- | --- |
|  | **Repeatability (CV^1^%)** | **Intermediate precision (CV%)** |
| AP2B1 - IQPGNPNYTLSLK | 10.2 | 18.3 |
| Gamma-synuclein - ENVVQSVTSVAEK | 26.4 | 26.4 |
| Neurogranin - KGPGPGGPGGAGVAR | 13.5 | 16.5 |
| NPTXR - NNYMYAR | 4.4 | 4.8 |
| NPTXR - LVEAFGGATK | 17.4 | 26.1 |
| NPTX1 - ETVLQQK | 9.0 | 9.0 |
| NPTX1 - CESQSTLDPGAGEAR | 4.5 | 8.5 |
| NPTX2 - VAELEDEK | 5.4 | 8.5 |
| NPTX2 - ETVVQQK | 10.3 | 10.3 |
| GDi-1 - QLICDPSYIPDR | 7.9 | 10.0 |
| PeBP-1 - NRPTSISWDGLDSGK | 15.2 | 17.7 |
| 14-3-3 zeta/delta - VVSSIEQK | 2.8 | 2.8 |
| syntaxin-1B - QHSAILAAPNPDEK | 24.2 | 24.2 |
| syntaxin-7 - EFGSLPTTPSEQR | 14.2 | 24.4 |

Notes: Data presented as CV (%). Abbreviations: ^1^Coefficient of variation.

| **Without Partial Volume Correction** | | | | | | | | | |
| --- | --- | --- | --- | --- | --- | --- | --- | --- | --- |
| **Synaptic Protein** | **ALL** | | | **CN^1^** | | | **AD^2^** | | |
|  | ***r*** | **n** | **p-value** | ***r*** | **n** | **p-value** | ***r*** | **n** | **p-value** |
| **AP2B1 - IQPGNPNYTLSLK** | -0.41 | 28 | 0.031* | -0.11 | 7 | 0.82 | -0.35 | 21 | 0.12 |
| **Gamma-synuclein - ENVVQSVTSVAEK** | -0.40 | 28 | 0.034* | -0.017 | 7 | 0.97 | -0.34 | 21 | 0.14 |
| **Neurogranin - KGPGPGGPGGAGVAR** | -0.41 | 28 | 0.032* | -0.39 | 7 | 0.39 | -0.27 | 21 | 0.23 |
| **NPTXR - NNYMYAR** | -0.31 | 27 | 0.11 | -0.28 | 7 | 0.54 | -0.25 | 21 | 0.28 |
| **NPTX1 - ETVLQQK** | -0.36 | 28 | 0.064 | -0.35 | 7 | 0.45 | -0.25 | 21 | 0.27 |
| **NPTX2 - VAELEDEK** | -0.11 | 27 | 0.57 | -0.32 | 7 | 0.48 | 0.083 | 21 | 0.72 |
| **GDI-1 - QLICDPSYIPDR** | -0.49 | 28 | 0.0082** | -0.29 | 7 | 0.53 | -0.34 | 21 | 0.084 |
| **PEBP-1 - NRPTSISWDGLDSGK** | -0.54 | 28 | 0.0029** | -0.29 | 7 | 0.53 | -0.47 | 21 | 0.033* |
| **14-3-3 zeta/delta - VVSSIEQK** | -0.33 | 28 | 0.086 | 0.19 | 7 | 0.69 | -0.28 | 21 | 0.23 |
| **syntaxin-1B - QHSAILAAPNPDEK** | -0.44 | 28 | 0.019* | 0.26 | 7 | 0.57 | -0.41 | 21 | 0.063 |
| **syntaxin-7 - EFGSLPTTPSEQR** | -0.44 | 28 | 0.019* | 0.26 | 7 | 0.58 | -0.49 | 21 | 0.024* |
| **With Partial Volume Correction** | | | | | | | | | |
| **Synaptic Protein** | **ALL** | | | **CN^1^** | | | **AD^2^** | | |
|  | ***r*** | **n** | **p-value** | ***r*** | **n** | **p-value** | ***r*** | **n** | **p-value** |
| **AP2B1 - IQPGNPNYTLSLK** | -0.40 | 28 | 0.04* | -0.10 | 7 | 0.83 | -0.38 | 21 | 0.09 |
| **Gamma-synuclein - ENVVQSVTSVAEK** | -0.35 | 28 | 0.06 | -0.04 | 7 | 0.93 | -0.33 | 21 | 0.15 |
| **Neurogranin - KGPGPGGPGGAGVAR** | -0.33 | 28 | 0.08 | -0.17 | 7 | 0.72 | -0.27 | 21 | 0.24 |
| **NPTXR - NNYMYAR** | -0.20 | 27 | 0.31 | -0.07 | 7 | 0.87 | -0.18 | 21 | 0.45 |
| **NPTX1 - ETVLQQK** | -0.24 | 28 | 0.21 | -0.14 | 7 | 0.77 | -0.20 | 21 | 0.40 |
| **NPTX2 - VAELEDEK** | 0.00 | 27 | 0.99 | -0.08 | 7 | 0.86 | 0.13 | 21 | 0.58 |
| **GDI-1 - QLICDPSYIPDR** | -0.42 | 28 | 0.02* | -0.25 | 7 | 0.59 | -0.38 | 21 | 0.09 |
| **PEBP-1 - NRPTSISWDGLDSGK** | -0.44 | 28 | 0.02* | -0.18 | 7 | 0.70 | -0.41 | 21 | 0.06 |
| **14-3-3 zeta/delta - VVSSIEQK** | -0.23 | 28 | 0.23 | 0.33 | 7 | 0.47 | -0.24 | 21 | 0.29 |
| **syntaxin-1B - QHSAILAAPNPDEK** | -0.35 | 28 | 0.07 | 0.36 | 7 | 0.43 | -0.37 | 21 | 0.10 |
| **syntaxin-7 - EFGSLPTTPSEQR** | -0.43 | 28 | 0.02* | 0.28 | 7 | 0.54 | -0.51 | 21 | 0.02* |

**Supplementary Table 3**. Associations between synaptic protein levels and global synaptic density (*DVR*) in a composite of AD-affected regions in the total sample as well as in the separate groups of Alzheimer’s disease and congitivly normal participants.

Notes: Associations between synaptic density and biomarker levels were explored with Pearson’s correlation analysis. Abbreviations: ^1^Cognitively Normal, ^2^Alzheimer’s disease; *p < 0.05
**Supplementary Table 4**. Associations between synaptic protein levels and hippocampal synaptic density in the total sample as well as in the separate groups of Alzheimer’s disease and congitivly normal participants.

|  | **Without Partial Volume Correction** | | |  | | |  | | |
| --- | --- | --- | --- | --- | --- | --- | --- | --- | --- |
| **Synaptic Protein** | **ALL** | | | **CN^1^** | | | **AD^2^** | | |
|  | ***r*** | **n** | **p-value** | ***r*** | **n** | **p-value** | ***r*** | **n** | **p-value** |
| **AP2B1 - IQPGNPNYTLSLK** | -0.013 | 28 | 0.95 | 0.049 | 7 | 0.92 | 0.18 | 21 | 0.45 |
| **Gamma-synuclein - ENVVQSVTSVAEK** | -0.065 | 28 | 0.74 | 0.30 | 7 | 0.51 | 0.11 | 21 | 0.62 |
| **Neurogranin - KGPGPGGPGGAGVAR** | 0.046 | 28 | 0.82 | 0.24 | 7 | 0.60 | 0.29 | 21 | 0.21 |
| **NPTXR - NNYMYAR** | 0.022 | 27 | 0.91 | 0.40 | 7 | 0.37 | 0.064 | 21 | 0.78 |
| **NPTX1 - ETVLQQK** | -0.0071 | 28 | 0.97 | 0.37 | 7 | 0.41 | 0.093 | 21 | 0.69 |
| **NPTX2 - VAELEDEK** | 0.175 | 27 | 0.38 | 0.41 | 7 | 0.36 | 0.31 | 21 | 0.17 |
| **GDI-1 - QLICDPSYIPDR** | -0.098 | 28 | 0.62 | 0.13 | 7 | 0.79 | 0.14 | 21 | 0.53 |
| **PEBP-1 - NRPTSISWDGLDSGK** | -0.087 | 28 | 0.66 | 0.15 | 7 | 0.75 | 0.17 | 21 | 0.47 |
| **14-3-3 zeta/delta - VVSSIEQK** | -0.031 | 28 | 0.88 | 0.093 | 7 | 0.84 | 0.23 | 21 | 0.31 |
| **syntaxin-1B - QHSAILAAPNPDEK** | -0.16 | 28 | 0.41 | -0.39 | 7 | 0.39 | 0.099 | 21 | 0.67 |
| **syntaxin-7 - EFGSLPTTPSEQR** | -0.096 | 28 | 0.63 | -0.020 | 7 | 0.97 | 0.062 | 21 | 0.79 |
| **With Partial Volume Correction** | | | | | | | | | |
| **Synaptic Protein** | **ALL** | | | **CN^1^** | | | **AD^2^** | | |
|  | ***r*** | **n** | **p-value** | ***r*** | **n** | **p-value** | ***r*** | **n** | **p-value** |
| **AP2B1 - IQPGNPNYTLSLK** | -0.02 | 28 | 0.91 | 0.01 | 7 | 0.99 | 0.17 | 21 | 0.46 |
| **Gamma-synuclein - ENVVQSVTSVAEK** | -0.08 | 28 | 0.69 | 0.26 | 7 | 0.58 | 0.10 | 21 | 0.65 |
| **Neurogranin - KGPGPGGPGGAGVAR** | 0.03 | 28 | 0.87 | 0.23 | 7 | 0.62 | 0.27 | 21 | 0.23 |
| **NPTXR - NNYMYAR** | 0.02 | 27 | 0.92 | 0.40 | 7 | 0.38 | 0.06 | 21 | 0.78 |
| **NPTX1 - ETVLQQK** | 0.00 | 28 | 0.98 | 0.37 | 7 | 0.42 | 0.10 | 21 | 0.67 |
| **NPTX2 - VAELEDEK** | 0.17 | 27 | 0.40 | 0.41 | 7 | 0.36 | 0.30 | 21 | 0.18 |
| **GDI-1 - QLICDPSYIPDR** | -0.11 | 28 | 0.58 | 0.09 | 7 | 0.84 | 0.14 | 21 | 0.55 |
| **PEBP-1 - NRPTSISWDGLDSGK** | -0.09 | 28 | 0.65 | 0.12 | 7 | 0.79 | 0.17 | 21 | 0.46 |
| **14-3-3 zeta/delta - VVSSIEQK** | -0.04 | 28 | 0.84 | 0.08 | 7 | 0.86 | 0.22 | 21 | 0.33 |
| **syntaxin-1B - QHSAILAAPNPDEK** | -0.17 | 28 | 0.38 | -0.41 | 7 | 0.36 | 0.09 | 21 | 0.70 |
| **syntaxin-7 - EFGSLPTTPSEQR** | -0.12 | 28 | 0.55 | -0.04 | 7 | 0.93 | 0.04 | 21 | 0.87 |

Notes: Associations between synaptic density and biomarker levels were explored with Pearson’s correlation analysis. Abbreviations: ^1^Cognitively Normal, ^2^Alzheimer’s disease
